# Development of Targetable Multi-Drug Nanoparticles for Glioblastoma Treatment and In Vitro Evaluation in Glioblastoma Stem Cells

**DOI:** 10.1101/2020.11.08.373696

**Authors:** Shelby B. Smiley, Yeonhee Yun, Pranav Ayyagari, Harlan E. Shannon, Karen E. Pollok, Michael W. Vannier, Sudip K. Das, Michael C. Veronesi

**Affiliations:** IUPUI; IU School of Medicine; University of Chicago Hospitals; Butler University

**Author notes:** Corresponding author and PI.

## Abstract

Glioblastoma (GBM) is a malignant brain tumor with a poor long-term prognosis. The current median survival is approximately fifteen to twenty months with the standard of care therapy which includes surgery, radiation, and chemotherapy. An important factor contributing to recurrence of GBM is high resistance of GBM cancer stem cells (CSCs) to several anticancer drugs, for which a systemically delivered single drug approach will be unlikely to produce a viable cure. Therefore, multidrug therapies have the potential to improve the survival time. Currently, only temozolomide (TMZ), which is a DNA alkylator, affects overall survival in GBM patients. CSCs regenerate rapidly and over-express a methyl transferase which overrides the DNA-alkylating mechanism of TMZ, leading to drug resistance. Idasanutlin (RG7388, R05503781) is a potent, selective MDM2 antagonist that additively kills GBM CSCs when combined with TMZ. Nanotechnology is an emerging field that shows great promise in drug delivery and diagnostics. The ability to combine both therapy and imaging allows real time assessment of drug delivery in vivo for the field of theranostics.

To develop a multi-drug therapy using multi-functional nanoparticles (NPs) that preferentially target the GBM CSC subpopulation and provide in vivo preclinical imaging capability. Polymer-micellar NPs composed of poly(styrene-*b*-ethylene oxide) (PS-*b*-PEO) and poly(lactic-*co*-glycolic) acid (PLGA) were developed investigating both single and double emulsion fabrication techniques as well as combinations of TMZ and RG7388. The NPs were covalently bound to a 15-base-pair CD133 aptamer in order to target the CD133 antigen expressed on the surface of GBM CSC subpopulation. For theranostic functionality, the NPs were also labelled with a radiotracer, Zirconium-89 (89Zr). The NPs maintained a small size of less than 100 nm, a low negative charge and exhibited the ability to effectively target and kill the CSC subpopulation. In addition, the conjugation of the CD133 aptamer was able to promote killing in CSCs leading to the justification of a targeted nanosystem to potentially improve localized therapy in future in vivo models. This work has provided a potentially therapeutic option for GBM specific for CSC targeting and theranostic imaging.

## Introduction

Glioblastoma multiforme (GBM) is a primary central nervous system disease with limited therapy options, a median survival period of approximately 15-20 months in the presence of aggressive treatment, and poor long-term outcome (1–3). Glioblastoma (GB) tumors exhibit heterogeneity of neuronal precursors, including abnormal astrocytic cell formation and are the most angiogenic brain tumor, displaying a high degree of endothelial cell hyperplasia and vascular proliferation (4). Standard of care therapies include surgery to remove the tumor, followed by chemotherapy drugs, and radiation therapy. Tumor treating fields are also being routinely applied given efficacy in a clinical trial (5). Tumor removal poses a vast challenge due to the tumor’s infiltrative nature and increased cellular heterogeneity making complete removal a remote possibility (6).

The high rate of GB recurrence is thought at least in part to be due to presence of a drug resistant cancer stem cell population (CSCs) (7). CSCs can readily generate both proliferating progenitor-like and differentiated tumor cells amid microenvironment cues (8). CSCs have increased DNA damage response systems (DDR) and reduced apoptotic mechanisms making them very resistant to current chemotherapy treatment (9). Although CSCs pose a great problem in TMZ resistance for GB, they may also provide an answer. CD133 is an important cell surface marker for CSCs in a variety of solid cancers, including those of the brain, prostate, pancreas, melanoma, colon, liver, lung and ovarian cancers (10). However, CD133 is also found on non-CSC tumor cells making sub-regional localization of CSC within the tumor difficult (11). Recent studies have shown the AC133 epitope within the CD133 more specifically targets the CSC population within a tumor (12).

Therefore, therapies that can preferentially target the CSC sub-population in GB by binding to the AC133 epitope on the CD133 cell surface protein are thought to have great potential to jump start progress in GB treatment (13). Given the complexity of working with antibodies, researchers have turned to aptamers against AC133 for their lack of immunogenicity, highly reproducible and stable conformation, and much smaller size (14). There is a critical and immediate need to develop novel multi-drug therapies that can eliminate the CSC population. Many single drug regimens delivered orally or intravenously are limited by difficulty with crossing the blood brain barrier (BBB) and dose-limiting peripheral side effects. To achieve better therapy, dual drug-loaded polymer-micellar nanoparticles (NPs) were prepared using the double emulsification solvent evaporation method and achieved a relatively uniform size of 85 nm. Drug combinations included current standard of care temozolomide (TMZ) in combination with RG7388. NPs were conjugated with anti-CD133 aptamer to aid in site-directed targeting to CSCs with 66% efficiency.

TMZ is a pro-drug that hydrolyzes to its active enzyme form at physiological pH and has proven effective at slowing or stopping cancer cell growth in the short run by acting as a DNA alkylating agent (15, 16). Although, TMZ is the most effective anti-neoplastic, chemotherapy drug and serves as the standard of care agent, the GB tumor usually recurs with one study reporting as many as 60% of patients with GB experience resistance to TMZ (17, 18). TMZ also has potentially harmful side effects, which include erratic DNA alkylation and cell death (15). TMZ has a short half-life of 1.8 hours necessitating multiple doses (19). Therefore, a single-drug treatment paradigm such as with TMZ can be ineffective because of the extensive heterogeneity of mutations and resistance weapons the tumor possesses.

It is suspected, that after standard of care treatment, the DDR is upregulated in the remaining cells. Therefore, addition of an inhibitor of the DDR could provide the additional treatment to diminish the remaining tumor. Small molecule murine double minute 2 (MDM2) inhibitor would prevent the repair of the DNA damage. MDM2 (known as HDM2 in humans) negatively regulates the tumor suppressor p53 by serving as an E3 ubiquitin ligase and is responsible for the export of p53 from the nucleus (17, 20, 21). Inhibition of MDM2 would reduce its oncogenic effects and could be therapeutic. Idasanutlin, RG7388, is of the nutlin class, is currently under clinical trials, is proven to bind specifically to MDM2, and has a >100-fold selectivity for GB in various cell lines (20). In addition, RG7388 has good systemic exposure, is metabolically stable in vivo, and is non-genotoxic (21, 22). A disadvantage to working with these inhibitors are the potential adverse effects to normal cells which would be improved by localized site directed delivery to the tumor before it can exert its mechanistic effects. Previous data from Chen et al has shown increased apoptosis for dual treatment of TMZ and RG7388 in neuroblastoma cell lines, which provides justifications for its use in the current study (23).

Multifunctional hybrid nanocarriers (MNCs) represent an emerging class of drug delivery vehicles that can allow the needed localized site directed therapy for the powerful one-two punch provided by TMZ and RG7388. MNCs have the potential to control the release of drug during both transport and at a specific location to reduce side effects (24). Packaging of the drugs in these NPs would create a protective “layer” between the drug and physiological environment to prevent premature alkylation and preserve its therapeutic activity. To construct the NPs, block copolymers polystyrene-*b*-polyethylene oxide (PS-*b*-PEO) and poly(lactic-*co*-glycolide) with or without functional capabilities were used to self-assemble into a micellar structure of NCs. The advantage of combining both the micellar and polymer components provide the opportunity for reduced size of the particle that could deliver the drug across blood brain barrier and allow to tune release kinetics while providing functionality. The NPs were synthesized via double emulsion-solvent evaporation method to be able to deliver both hydrophobic and hydrophilic drugs. In recent articles, TMZ has been encapsulated traditionally using a single emulsion technique. However, because of its limited solubility in hydrophobic solvents such as chloroform or dichloromethane (DCM), its encapsulation efficiency has been exceedingly poor (25, 26). TMZ is an amphiphilic drug and treating it as a hydrophilic component may increase encapsulation efficiency. RG7388 is a hydrophobic drug and through the double emulsion technique can be co-loaded into the NPs with TMZ.

Objective of the study was to develop CD133 aptamer and ^89^Zr conjugated, TMZ and RG7388 loaded polymer-micellar NPs of size range around 100 nM, with small charge that prevents aggregation, and study their physicochemical properties and cell killing efficacy in GBM cell lines.

## Materials and Methods

The materials used in this study and their suppliers are documented in the APPENDIX.

### Determination of 0.5% 13k polyvinyl alcohol viscosity

To determine the viscosity of 0.5% 13k PVA, the Bohlin CVO 100 Rheometer from Malvern Panalytical (Malvern, UK) was used. The viscosity of the solution was measured in triplicate and was collected as an average value of the instantaneous velocity once the instrument levelled off in values.

### Analysis of TMZ by UV-Vis spectroscopy

For analysis of NPs encapsulating multiple drugs, methods were developed to analyze drug content. To analyze TMZ by UV-Vis for the single emulsion NPs, absorbance was measured at 325 nm, comparable to the literature maximum of 328 nm (27). A standard curve was made in triplicate. To improve sensitivity of the detection by UV-Vis spectroscopy, prior to beginning double emulsion fabrication of NPs, the maximum wavelength was determined on the UV-Vis spectrophotometer. A scan from 270 nm to 380 nm was conducted to determine the maximum TMZ absorption wavelength in dimethyl sulfoxide (DMSO). A standard curve was generated reading at 332 nm and was prepared in triplicate to measure drug content.

### Analysis of polymers by UV-Vis spectroscopy

To determine if polymers interfere with UV-Vis analysis of TMZ content, absorbance scans of each polymer dissolved in DMSO was conducted.

### TMZ stability

Because TMZ hydrolyzes at physiologic pH, it was necessary to determine the stability of TMZ in aqueous solutions at various pH to improve the formulation of the NPs. TMZ was dissolved in deionized water at pH 3.48 (pH 4), pH 4.87 (pH 5), pH 6.88 (pH 7) and analyzed by UV-Vis spectroscopy scanning from 220 nm to 370 nm to monitor for degradation or the presence of new peaks as an indication of conversion to MTIC/AIC. Measurements were conducted in triplicate over the course of two weeks.

### Determination of TMZ and RG7388 by HPLC

High pressure liquid chromatography (HPLC) was used as a quantitative method for the determination of drug content on an Autosampler (Agilent 1200 Dual-Loop Series, Agilent Technologies, Santa Clara, CA). A method was developed for the separation of TMZ and RG7388. TMZ and RG7388 were dissolved in ACN and were separated using a mobile phase of water and ACN. The samples were injected through a Zorbax Eclipse C8 4.6 x 150 mm 5 *μm* column (Agilent Technologies, Santa Clara, CA). Reference wavelengths for all three methods were set to 332 nm for TMZ and 273 nm for RG7388. Between each drug component, a flat baseline was established to accurately measure concentration. The first method used to separate the two drugs was based on a 50:50 ratio of water and ACN. The injection volume was 3 *μ*L, flow rate was 1 mL/min and the temperature was 40 °C. The next method used to separate the drugs required solvent ratios shown in Table 3.2. However, the water was not mixed with formic acid initially and the flow rate was 1 mL/min with an injection volume of 3 *μ*L.

### Nanoparticle fabrication by single emulsion method

A solvent emulsion evaporation system was adapted from previous work of Nabar et al. to initially fabricate the NPs (28). To form the oil phase for TMZ-loaded NPs, 200 *μ*L of either 0.1% PS(5.0k)-*b*-PEO(2.2k) or 0.1% PS(9.5k)-*b*-PEO(18k) dissolved in DCM and 20 *μ*L of 5% 50:50 73k PLGA dissolved in DCM were combined with 500 *μ*L of 0.1% TMZ dissolved in 135 *μ*L of DMSO and 2.865 mL of DCM. The organic phase was vortexed for 30 seconds and added dropwise to 8 mL of 0.5% 13k PVA. During addition of the organic phase, the solution was sonicated using a Branson 250 probe sonicator from Branson Ultrasonics (Danbury, CT) at constant duty cycle for five minutes at 20% power over ice. After sonication, the emulsions were stirred at 650 rpm for 2.5 hours to allow the organic solvents to completely evaporate. Each sample, unless otherwise noted, was prepared in triplicate. TMZ+RG7388 NPs were fabricated using the same method, however, the 500 *μ*L of drug added contained 0.1% solution of TMZ and 0.1% RG7388 dissolved in 225 *μ*L of DMSO and 2.775 mL of DCM. Control NPs were fabricated the same way without the drugs added and consisted of empty PLGA-PS-*b*-PEO particles, PLGA particles, and PS-*b*-PEO particles. PS-*b*-PEO control micelles were fabricated using 800 *μL* of 0.1% of PS-*b-*PEO to improve polymer concentration.

A similar method to Nabar et al. was used and nanoparticles were filtered using a 0.45 *μ*m filter to remove any aggregates (28). NPs were then centrifuged at 2,300 rcf for one hour, supernatant removed and replaced with MilliQ water. This process was then repeated two more times for thirty-minute cycles each to remove adsorbed PVA and other highly water-soluble components. These low speeds were used for purification to prevent aggregation of the NPs (28).

### Nanoparticle fabrication by double emulsion method

A double emulsification solvent evaporation technique adapted from Xu et al. was used to prepare the NPs (29). Prior to choosing a final method for the double emulsion NPs, many combinations and techniques were tested including sonication time, molecular weight of PLGA, surfactant type, surfactant molecular weight, surfactant concentration, technique of transfer and organic to aqueous phase volume ratios. Table 1 outlines the different methods tested during the preliminary discovery stages. The size of the NPs were the determining factor in selecting the final method.

**Table 1:**
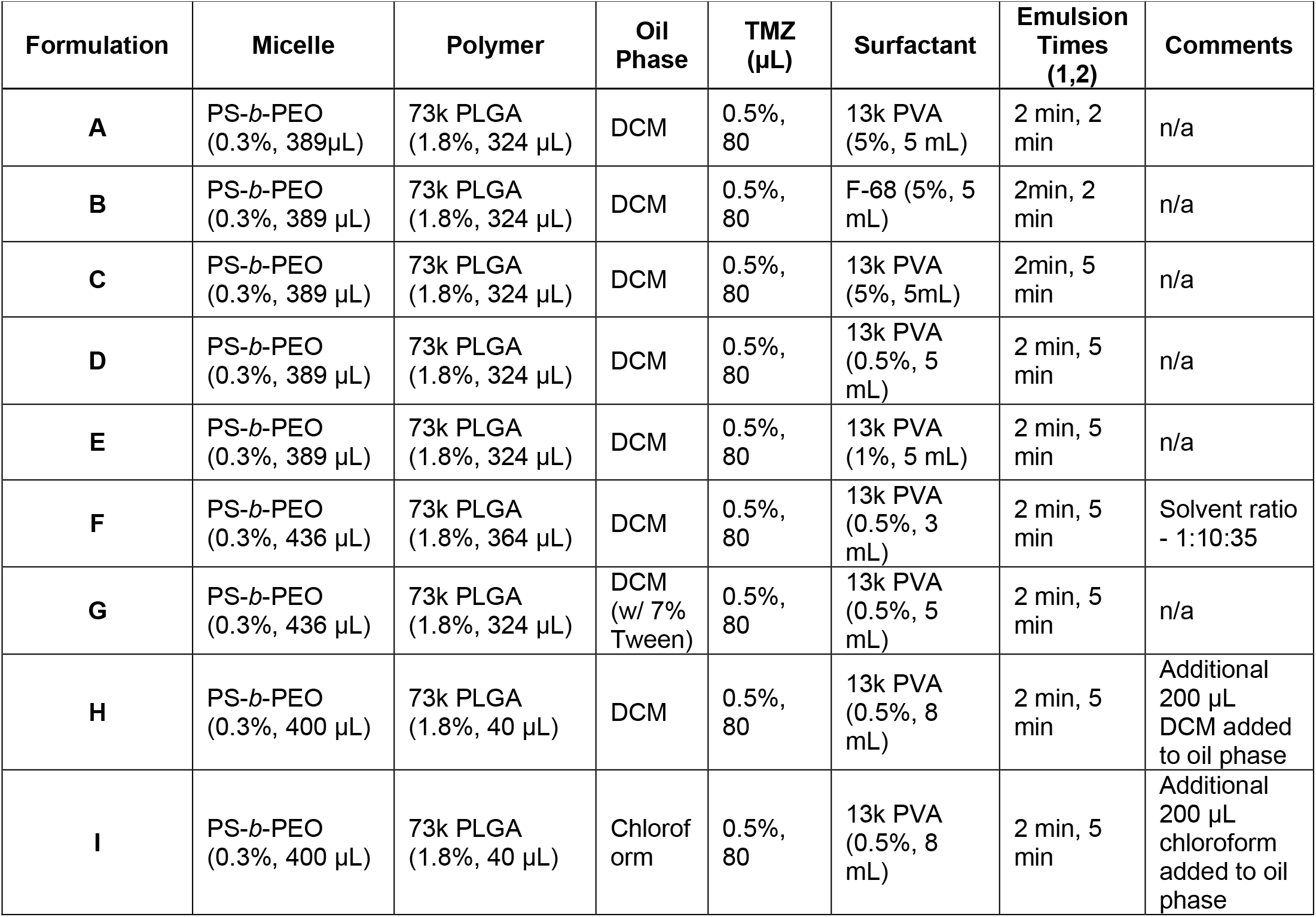
Double Emulsion Discovery Stages.

The method for double-emulsion NPs maintained the same PLGA:PS-*b*-PEO weight ratio of 1:5 as used in the single-emulsion NPs (28). To synthesize NPs with TMZ only, 389 μL of 0.3% PS(9.5k)-*b*-PEO(18k) was dissolved in DCM for at least 30 minutes and combined with either 324 μL of 1.8% 50:50 73k PLGA or 162 μL of 1.8% 50:50 50k PLGA-NHS and 162 μL of 1.8% 50:50 30k PLGA-NH2 dissolved in DCM for about 30 minutes. The organic phase was vortexed for 30 seconds. This organic phase was sonicated in ice bath using the Branson 250 probe sonicator at constant duty cycle for 2 minutes at 20% power. Immediately after starting the sonication, 80 μL of 0.4% TMZ in 0.1N HCl, which had been thoroughly dissolved using bath sonication and applying heat, was added dropwise to form the first emulsion. Once completed, the first emulsion was added dropwise to 4 mL 0.5% 13k PVA at pH 4 for a second emulsion at constant duty cycle for 5 minutes at 20% power. An additional 1 mL of PVA was used to wash the remaining particles from the first emulsion into the second emulsion. After sonication, the NPs were left to stir for 2.5 hours at 650 rpm for evaporation of the organic solvents.

To fabricate dual-drug NPs, 43 μL of 0.2% RG7388 dissolved in ethyl acetate dissolved in DCM was added to the initial organic phase after the 30 second polymer vortex. The entire organic phase was then vortexed for an additional 30 seconds.

To synthesize NPs with TMZ only, PLGA-NHS (162 μL) at a fixed stock concentration of 1.8% was added to PLGA-NH2 (162 μL, 1.8%) and carboxylated PS-*b*-PEO (389 μL, 0.3%) in DCM. Solutions were vortexed for 30 seconds. TMZ (80 μL, 0.4%) was added dropwise to the oil phase during sonication. These biphasic solutions were sonicated using a probe sonicator (Branson Sonifier 250) at a constant duty cycle for 3 min. This solution was then immediately added dropwise to aqueous PVA (5 mL, 0.5%), at a pH of 4, during a second sonication at a constant duty cycle for 5 min. The resulting emulsion was left to stir for 2.5 hours at 650 rpm to allow evaporation of the organic solvents. For this method, the NPs were filtered using the 0.45 μm filter and then were collected using an Amicon Ultra-15 centrifugal filter from Millipore Sigma (Burlington, MA) with a 100 kDa molecular weight cut off. NPs were added to the filter and centrifuged at 5,000 rcf for 20 minutes. The NPs were washed with deionized water pH 4 twice for 15 minutes at 5,000 rcf and for a final wash and centrifugation of 40 minutes at 5,000 rcf. The remaining suspended NPs were then either used immediately or lyophilized in a Labconco freeze dry system, Freezone 4.5 from Labconco Corporation (Kansas City, MO) until further use.

### Nanoparticle characterization

#### Size, polydispersity index, and charge by dynamic light scattering

A Malvern Panalytical Zetasizer (Malvern Panalytical, Malvern, UK) was used to measure the hydrodynamic diameters of the particles through dynamic light scattering (DLS). Particles were dropped into either water or 0.5% 13k PVA with viscosity values set to 0.8872 cPs or 2.78 cPs, respectively. The viscosity for 0.5% 13k PVA was used after the initial discovery stages of double emulsion particle fabrication. The particle concentration was adjusted to maintain a scattered light intensity between 100 and 200 kilocounts per second. Laser illumination was applied at 633 nm and the detector was set to 90^°^. This measurement determined the PDI to serve as a quantitative analysis of uniformity within the sample of NPs.

The Malvern Panalytical Zetasizer was also used to measure the zeta-potential or surface charge of the NPs. Laser illumination was applied at 633 nm and the detector was set to 90°. Particle size and zeta-potential were measured prior to collection.

#### Size and PDI stability of NPs

TMZ-loaded PS-*b*-PEO+PLGA-NHS/PLGA-NH2 NPs were measured at various time points in order to assess size and PDI stability when the NPs are stored at 4°C in the 0.5% 13k PVA solution at a pH≤4. Size and PDI were measured with DLS at time zero, 24 hours, and 8 days.

### Transmission electron microscopy

NPs were placed on a formvar/carbon-coated 200 mesh grid (Electron Microscopy Sciences, Hatfield, PA) and allowed to absorb for three minutes after which the remaining solution was wicked away. The grid was then placed on a drop of 1% uranyl acetate in distilled water for three minutes. The grid was allowed to dry on filter paper. It was then viewed on a Transmission Electron Microscope (ThermoFisher Spirit, Hillsboro, OR), and images were taken with a CCD Camera (Advanced Microscopy Techniques, Danvers, MA).

### Drug encapsulation and drug-loading percentage by UV-Vis spectroscopy

The TMZ-loaded NPs were fabricated using the same methods, however, for single emulsion NPs, five times the initial concentration of TMZ stock was used (0.5% instead of 0.1%). The evaporated NPs for single and double emulsion were purified to remove un-encapsulated TMZ using their respective methods. The washed pellets were dried under vacuum. For measurement of single emulsion NPs, the pellets were dissolved into a known volume of DMSO and were measured according to a literature value for the maximum absorbance of TMZ, 325 nm, and were compared to a standard curve. For measurement of double emulsion NPs, additional steps were taken to remove absorption interference from PS-*b*-PEO. The pellets were dissolved in DMSO to release the encapsulated TMZ into solution. Then, the PS-*b*-PEO was precipitated with 0.1N HCl and filtered with 0.22 μm filter.

TMZ was then measured by UV-Vis spectroscopy at 332 nm and compared to a standard curve. Drug encapsulation efficiency was determined by the following equation:

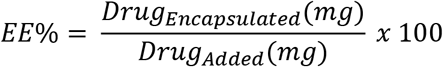

Drug loading percentage was determined by the following equation:

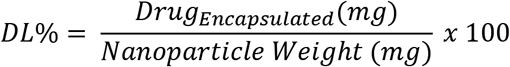

### Drug encapsulation and drug-loading percentage by HPLC

A standard curve of either TMZ or both TMZ and RG7388 dissolved in ACN was measured with HPLC on the same day as the analysis to avoid any environmental changes from day to day measurements. The standard curve was made starting with 54 μg/mL of TMZ and 51.6 μg/mL of RG7388 and diluted to make at least five standards. These were plotted by comparing concentration to area under the curve.

Standards and unknowns were injected at a volume of 100 μL through a Zorbax Eclipse C8 4.6 x 150 mm 5 μm column. Mobile phase consisted of ACN and water adjusted to pH 4 using formic acid and the samples were analyzed according to the following gradient method in Table 2.

**Table 2:** Method for HPLC separation of TMZ and RG7388

| Time (min) | Water (%) | ACN (%) | Flow (mL/min) |
| --- | --- | --- | --- |
| 0.0 | 20.0 | 80.0 | 0.8 |
| 2.0 | 20.0 | 80.0 | 0.8 |
| 7.0 | 5.0 | 95.0 | 0.8 |
| 10.0 | 0.0 | 100.0 | 0.8 |
| 12.0 | 0.0 | 100.0 | 0.8 |
| 13.0 | 20.0 | 80.0 | 0.8 |

### Conjugation of anti-CD133 aptamer

To conjugate the aptamer to the NPs containing PLGA-NHS, the NHS to amine reaction was employed following click chemistry. Approximately 7 mg of NPs were suspended in DNAse and RNAse free PBS at pH 5. 20 μL of the anti-CD133 aptamer, either fluorescent or non-fluorescent, was added from a stock solution of 100 μM. Nanoparticles were then protected from light and stirred at room temperature at 600 rpm for 2 hours. After stirring, the NPs were collected by ultra-centrifugation using a Centriprep-10 with a molecular weight cut off of 10 kDa for 20 minutes at 3,000 rcf. The filtrate contained free, unbound aptamer while the substrate contained the NPs bound to the aptamer. The NPs were then washed using DNAse and RNAse free water and ultra-centrifuged for 15 minutes at 3,000 g.

To measure the efficiency of the aptamer, the fluorescent anti-CD133 aptamer with the FAM linker was conjugated. A standard curve consisting of six known standards was made and measured each time using a Victor 3V 1420 Multilabel Counter from Perkin Elmer (Waltham, MA). Fluorescence was measured by using an excitation wavelength of 485 nm and an emission wavelength of 535 nm. The fluorescence of the aptamer bound to the NPs was then compared to the standard curve. Binding efficiency was determined by the following equation:

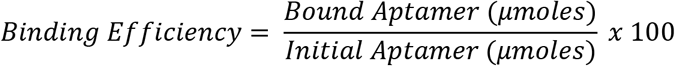

### Stability of anti-CD133 aptamer

Preliminary studies were conducted to analyze the stability of the aptamer bound to the NPs. To analyze whether the aptamer was stably bound to the NPs, an electrophoretic mobility shift assay (EMSA) was conducted. NPs were conjugated to the anti-CD133 aptamer. NPs were ran on a horizontal 1.5% agarose gel for forty minutes at 150 V in 1X TAE buffer. Lane 1 consisted of 10 μL from a stock solution of trypan blue (10 μL), DNAse/RNAse free water (90 μL), and glycerol (2 μL). Lane 2 contained 10 μL of a stock solution containing free fluorescent aptamer (1 μL), glycerol (20 μL), and DNAse/RNAse free water (179 μL). Lanes 3-5 contained different batches of conjugated NPs. These lanes were loaded with 10 μL of solution from a stock that contained 10 μL of aptamer bound NPs and 2 μL of glycerol. The gel system was placed in ice and protected from light.

To analyze the binding stability over time, the binding efficiency of the aptamer was measured using the same method as above. The sample was then placed in the 37^°^C incubator to mimic in vitro conditions. At various time points, the NPs were centrifuged in a Centriprep-10 centrifugal filter with a molecular weight cut off of 10 kDa for 15 minutes at 3,000 g. The time points were 0 hours, 0.5 hours, 1 hour, 3 hours, and 24 hours. A known volume of NPs with the bound aptamer was collected and compared against a standard curve by analyzing with the plate reader under the same conditions previously listed. Due to constraints of time available in the lab, this experiment was conducted with a sample size of one.

### Conjugation of ^89^Zr

To conjugate ^89^Zr to the NPs, two different methods were explored for determining an optimum binding efficiency. First, 1.5 mg of aptamer-conjugated NPs were dispersed in 2 mL of HEPES buffer to neutralize any acidity. The NPs used were washed with HEPES buffer after conjugation to the aptamer. This was then added to 1 mL of 20 mM deferoxamine mesylate (DFOM) in water, the mesylate salt of deferoxamine (DFO). The NPs were then stirred for one hour at room temperature to conjugate the amine group and DFOM. Excess DFOM was then removed by ultra-centrifugation for 40 minutes at 5,000 g using an Amicon Ultra-15 with a molecular weight cut off of 100 kDa. NPs were washed with HEPES buffer for an additional spin of 40 minutes at 5,000 g. ^89^Zr oxalate was neutralized using 60 μL of HEPES buffer and 60 μL of potassium carbonate (K_2_CO_3_). 200 μCi of ^89^Zr was added to the NPs, mixed and then incubated for 1 hour at 37°C. NPs were then added to HEPES buffer and centrifuged under the same conditions. This method was conducted in triplicate.

The second method was similar with slight modifications. An entire batch of aptamer-conjugated NPs containing approximately 7 mg of NPs were washed with HEPES buffer and suspended in 2 mL of 20 mM DFOM in water. This solution was stirred overnight at room temperature to conjugate the amine group to the DFOM. NPs were collected after adding the solution to 10 mL of HEPES buffer with the same method. NPs were collected from the filter using 7 mL of HEPES buffer and 0.5 mL of the solution, containing approximately 0.5 mg of NPs, was mixed with 1 mCi of ^89^Zr and incubated for 2 hours at 37°C. NPs were then added to HEPES buffer to neutralize the reaction and were centrifuged under the same conditions. This method was conducted in triplicate.

### Binding efficiency and stability of ^89^Zr

To determine the binding efficiency of the ^89^Zr to the NPs, the radioactivity was measured using a dose calibrator from Biodex Medical Systems (Shirley, NY). The radioactivity of the pellet of NPs, the NPs remaining in initial vial, and the supernatant were all measured to account for all traces of ^89^Zr. The efficiency was determined by importing the radioactivity of the following samples:

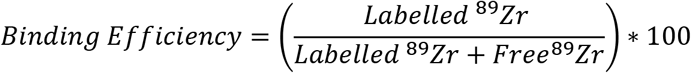

Stability of ^89^Zr bound to the NPs was only conducted using the first method. After measuring binding efficiency, the pellet of NPs was added to HEPES buffer and allowed to stir at 37 °C for 24 hours. The NPs were then collected using the ultracentrifugation method previously described and the radioactivity was measured. To account for radioactive decay, the radioactivity of the entire solution was measured prior to collection.

### In vitro studies in glioblastoma cell culture

Human GBM43 cells were kindly donated from the Simon Cancer Center at Indiana University. Culture media was Gibco Dulbecco’s Modified Eagle Medium (DMEM) with 4.5 g/L D-Glucose and L-Glutamine with 10% Fetal bovine serum (FBS) and 1% HEPES buffer (1M pH 7.3). These cells could be seeded for testing in 96-well plates at approximately 85%-90% confluency. GBM CSC’s were cultured using the Human Glioma Cancer Stem Cells Media with serum requested without antibiotics approximately 60% to 80%confluency according to the Celprogen Human Glioblastoma Cancer Stem Cell seeding protocol. Both cell lines were incubated at 37^°^C in a humidified atmosphere containing 5% carbon dioxide (CO_2_).

### Determination of optimal cancer stem cell seeding number

Prior to performing cytotoxicity studies of both free drugs and NPs, a cell seeding number for the study in 96-well plates was first optimized. To determine the optimal cell seeding number for CSCs, cells were treated with RG7388. Cells were plated in triplicate at concentrations of 100 cells/well, 50 cells/well, and 25 cells/well and left to culture overnight. Cells were then treated with RG7388 at concentrations of 50 μM, 37.5 μM, 28.13 μM, 21.09 μM, 15.82 μM, 11.87 μM, 8.90 μM, and 6.67 μM. Cytotoxicity was analyzed by the methylene blue assay listed in the following subsection. The dose curves were then compared to determine the optimal cell seeding number.

### Determination of IC_50_ Values

To determine the IC_50_ value for each drug, a methylene blue assay determined the cytotoxicity at varying doses. GBM CSCs were plated in a 96-well plate at 25 cells/well in a volume of 100 μL and cultured overnight. Media was adjusted to a series of single drug concentrations as seen in Table 3. A vehicle treatment was also used that obtained the maximum DMSO concentration used. DMSO concentration was maintained ≤0.1% of the total media concentration. In addition, studies were conducted with media controls on each plate.

**Table 3:** Single drug doses of the drugs.

| Cell Line | Drug | Doses |  |  |  |  |  |  |  |
| --- | --- | --- | --- | --- | --- | --- | --- | --- | --- |
| GBM43 | TMZ (μM) | 3400 | 1700 | 850 | 425 | 212.5 | 106.3 | 56.1 | 26.6 |
| CSCs | TMZ (μM) | 2000 | 1000 | 500 | 250 | 125 | 62.5 | 31.3 | 15.6 |
|  | RG7388 (μM) | 40 | 30 | 22.5 | 16.9 | 12.7 | 9.5 | 7.1 | 5.3 |

Combination treatments were conducted next. TMZ and RG7388 were combined for CSC treatment at ratios of 15:1, 50:1 and 500:1 starting with TMZ concentrations of 600 μM, 500 μM and 500 μM, respectively. Treatment groups of NPs were empty non-targeted NPs, non-targeted TMZ-loaded NPs, non-targeted TMZ+RG7388-loaded NPs, and targeted TMZ+RG7388loaded NPs in CSCs. The NPs used for in vitro studies were the final formulation of NPs seen in Table 1 The highest treatment for each of the four groups contained 714 μg of NPs and were diluted by half for a total of nine treatments.

Cells were incubated with the treatments for five days. After, the media was aspirated, and the cells were fixed with 100% methanol for 5 to 10 minutes. Methanol was disposed of and the wells were allowed to dry completely. The remaining viable cells were stained with 70 μL 0.05% methylene blue for 30 to 60 minutes. The plates were then shaken into the sink, dipped completely into a pitcher of deionized water to rinse three times, and the water was flicked out. The remaining water was shaken out onto a paper towel and the plates were allowed to dry completely. Next, 100 μL 0.5M HCl was added using a multi-channel pipette. The plates were mixed by gently tapping and read at 600 nm on the plate reader described above. Treatment amounts will be listed with the results, but goal treatment duration was three trials in three different cell populations to account for population differences.

To determine the cytotoxicity, Fa, the fraction of cell affected by the drug, and Fu, the fraction unaffected by the drug, were determined by the following equations:

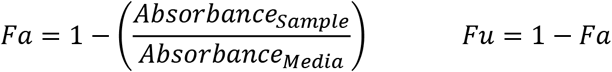

These values were then plotted and analyzed in CalcuSyn 2.0 to determine an IC_50_ value.

### Statistics

Unless otherwise stated, experiments were conducted in at least triplicate and a standard error of the mean was used. Statistics were conducted with a confidence value of 95%. A students t-test was used to conduct comparison between groups. All data was analyzed using a two-tailed, unpaired analysis with the assumption of unequal variances. The only test that included a paired analysis was that of the size stability of NPs. For each sample, data is reported as mean size ± standard error of the mean.

## Results

### Particle constituent analysis

Viscosity was measured by averaging the values of the instantaneous viscosity after the instrument levelled off at a value for viscosity. These levelled values can be seen Figure 1. An average of the three trials produced a viscosity measurement of 1.59*10^-7^ Pa*s or 2.788±10^-5^ cPs.

**Figure 1:**
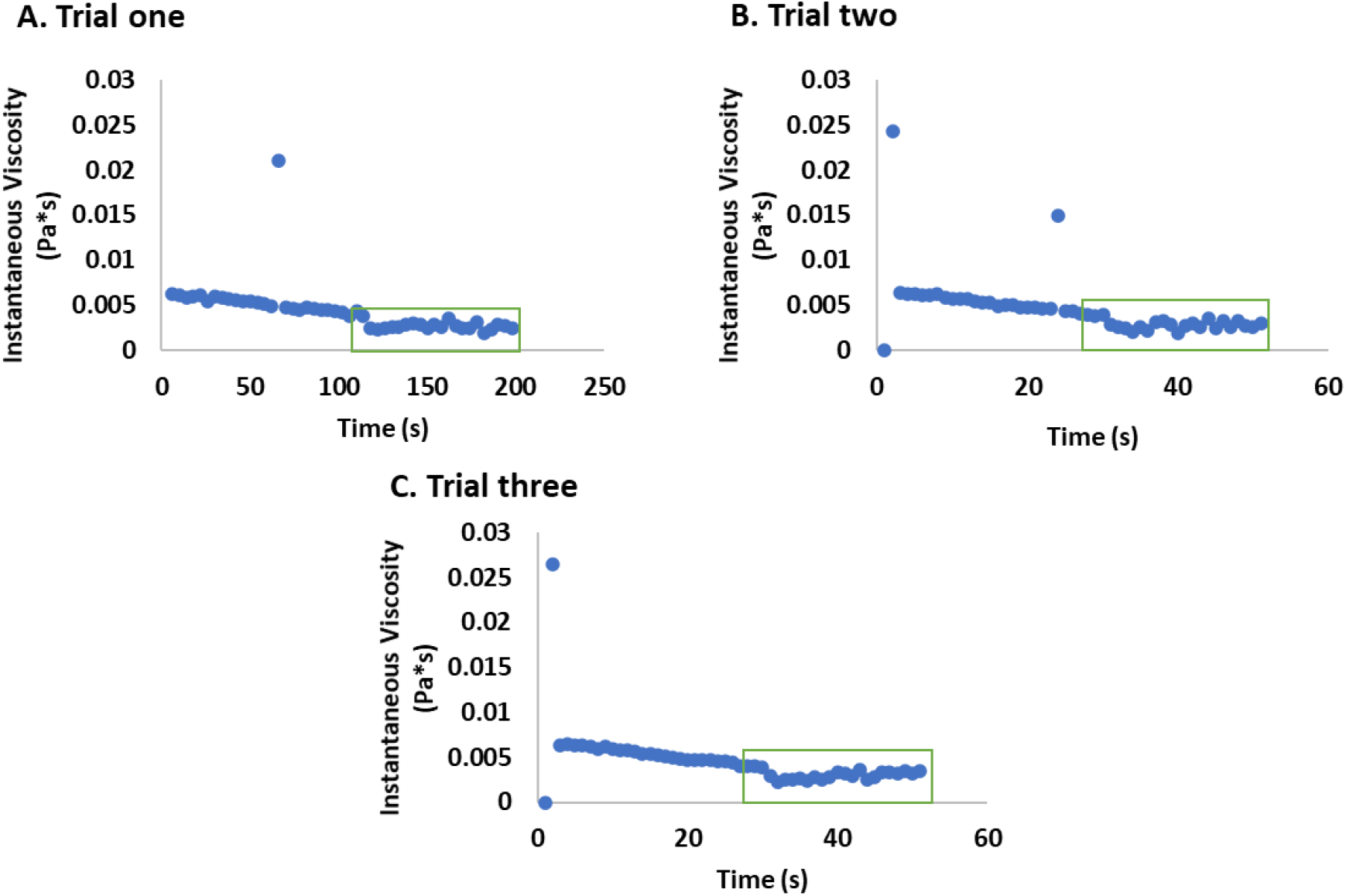
Determination of surfactant viscosity. Above represents the three trials of the measurement of viscosity for 0.5% 13k PVA with a rheometer. The data set that is used in viscosity determination is boxed in green.

To determine if the polymers interfere with analysis by UV-Vis spectroscopy at 332 nm, absorbance scans were measured. It was determined from Figure 2 that PS-*b*-PEO has absorbance around the region similar to TMZ, while PLGA does not.

**Figure 2:**
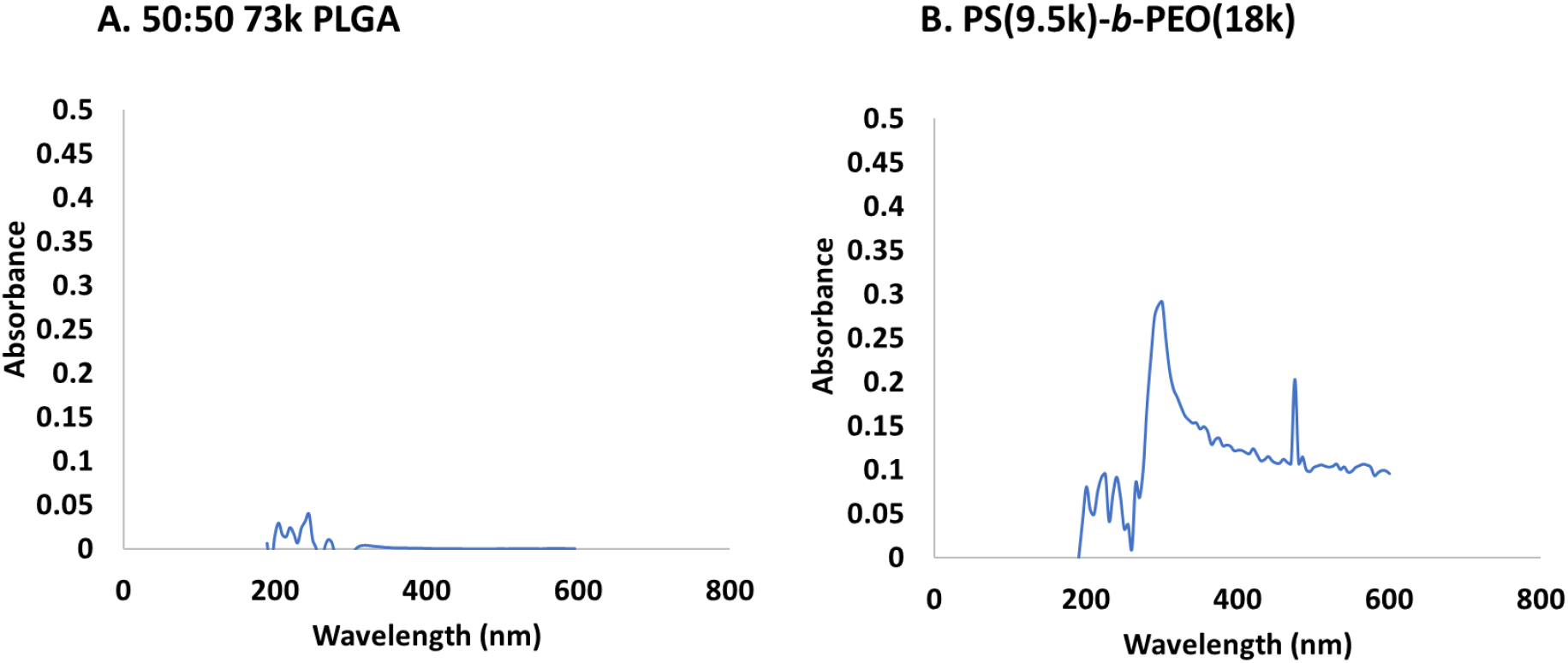
Polymer absorbance scans in DMSO. These polymer scans were conducted by UV-Vis spectroscopy to determine if either 50:50 73k PLGA or PS(9.5)-b-PEO(18k) have absorbance in the same regions as TMZ.

Scans of TMZ at various pH of aqueous solutions were analyzed by UV-Vis over time. These scans can be seen in Figure 3. The slopes from these curves in order of increasing pH are −0.0037 nm per day and −0.0077 per day.

**Figure 3:**
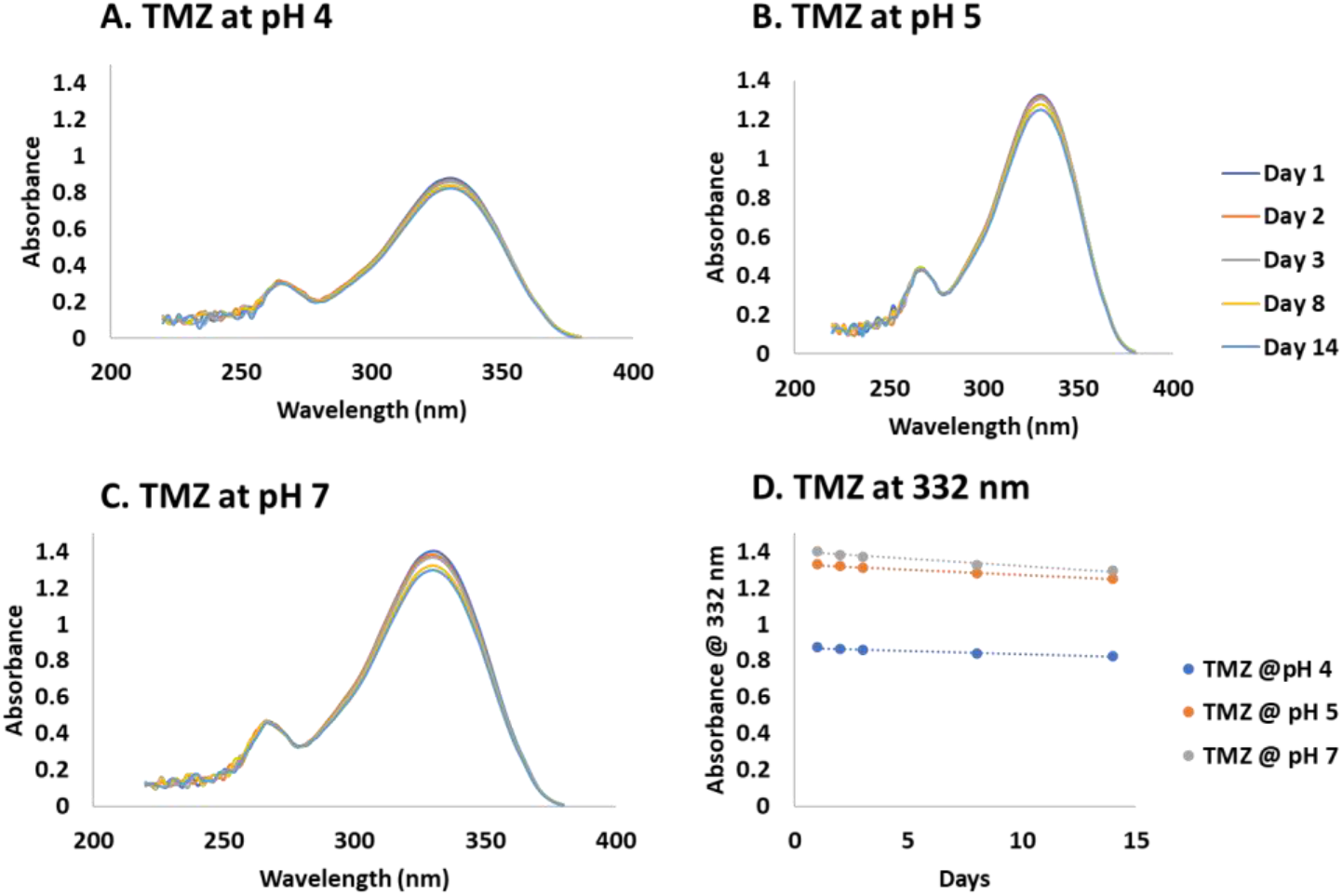
Stability of TMZ. The stability of TMZ was measured at at pH 4 (A), pH 5 (B), pH7 (C), as well as a linear regression at maximum absorbance of 332 nm (D).

### Method determination for HPLC drug analysis

Two different methods were investigated to separate TMZ from RG7388. For the first isocratic method, the release times for the TMZ and RG7388 were approximately 3.0 min and 1.81 min, respectively. The release times for the drugs using the second method were approximately 3.09 and 6.31 min. The final method used for drug quantification shorted the release times of TMZ and RG7388 to 1.9 min and 3.8 min while a baseline was established with a much broader peak. Reference wavelengths of 332 nm and 273 nm were used for TMZ and RG7388, respectively.

### Effect of the method of preparation on particle size, morphology, and surface charge

Size and PDI were measured by DLS and are reported in Table 4. These results are the sizes and PDI values that are associated with the initial formulation development stages for double emulsion NPs corresponding to the particle formulation described in Table 1. Size was determined by an average volume distribution. Because the viscosity value of water was used during measurement of DLS, these results are unable to be compared statistically or to the final formulation. However, sizes between the groups can be compared. It was determined that increasing the concentration of the PVA surfactant, results in a decrease in hydrodynamic diameter and reduction of PDI. In addition, swapping PVA for F-68 resulted in an increase in particle size. An increase in sonication time slightly reduces the size of NPs but might not have a great effect. In addition, the use of Tween in the organic layer did not reduce the size of the particles but reduced the PDI.

**Table 4:** Size and PDI values of initial formulation of double-emulsion particles

| Particle Description | Size (nm) | PDI |
| --- | --- | --- |
| A | 195±9 | 0.103 |
| B | 380±4 | 0.111 |
| C | 142±7 | 0.179 |
| D | 141±15 | 0.086 |
| E | 263±6 | 0.128 |
| F | 204±6 | 0.128 |
| G | 210±4 | 0.108 |
| H | 225±11 | 0.152 |
| I | 210±11 | 0.211 |
| J | 207±2 | 0.161 |
| K | 169±13 | 0.185 |

A final formulation of NPs was selected by choosing the smallest size and a size that was less than 100 nm set by the initial size objective. These results were conducted after the initial formulation development in the preceding paragraph and were used for the rest of the article. Characteristics of these NPs can be seen in Table 5 to accomplish the first objective. Empty polymer-micellar NPs had a size of 95 nm and there was a significant decrease in size to 82 nm upon the addition of TMZ. When RG7388 was loaded into the NPs, there was not a significant change in size. There was a significant decrease in zeta-potential from −7.1 mV to −9.9 mV when RG7388 was added into the TMZ-loaded NPs.

**Table 5:** Characteristics of final double emulsion NP formulation

| Particle Description | Size (nm) | PDI | Charge (mV) |
| --- | --- | --- | --- |
| PS-*b*-PEO+PLGA-NHS/PLGA-NH<sub>2</sub> | 95±3 | 0.126 | -8.0±1.0 |
| PS-*b*-PEO+PLGA-NHS/PLGA-NH<sub>2</sub>+TMZ | 82±3 | 0.175 | -7.1±0.4 |
| PS-*b*-PEO+PLGA-NHS/PLGA-NH<sub>2</sub>+TMZ+RG7388 | 87±2 | 0.138 | -9.9±0.7 |

TMZ-loaded PS-*b*-PEO+PLGA-NHS/PLGA-NH_2_ NPs were measured over time to determine size and PDI stability. Size increased over time, but there was only a significant increase of size between 0 and 24 hours (Figure 4). PDI did not show a trend and there was no significant change (Figure 5).

**Figure 4:**
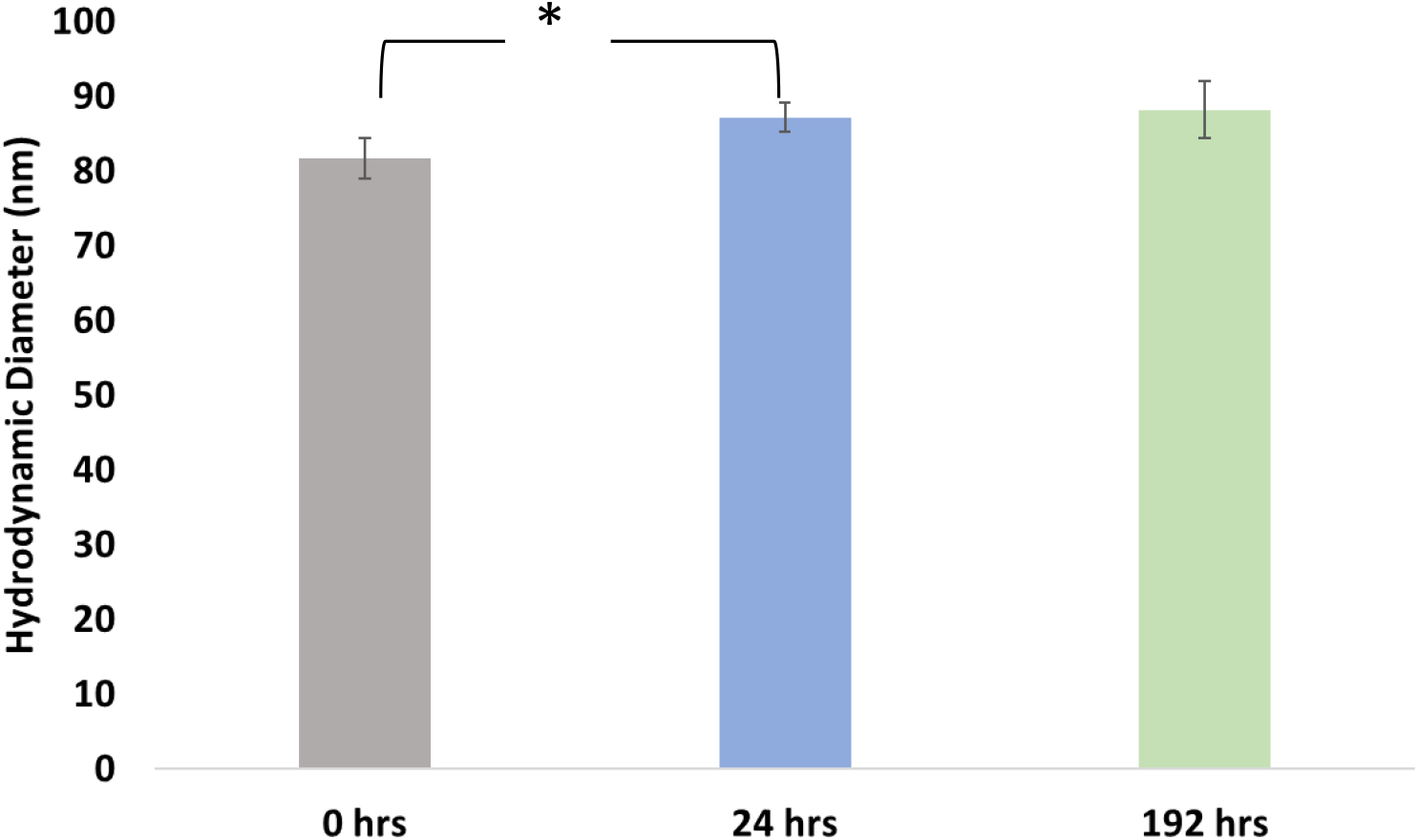
Size stability of NPs over time. Size of NPs were measured at each time point. Each bar corresponds with the PDI results seen in Figure 5. Significance is represented by an (*).

**Figure 5:**
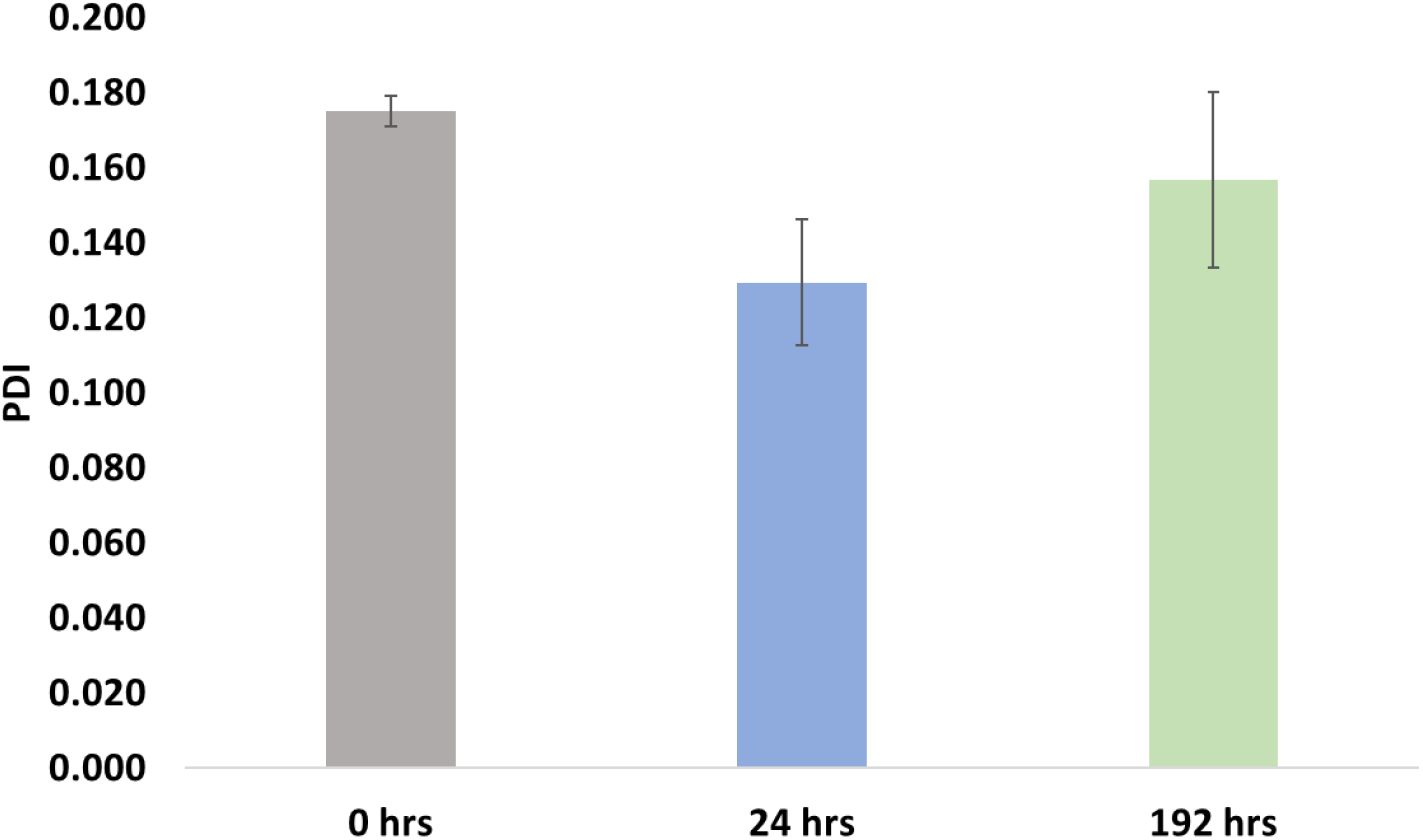
PDI stability of NPs over time. PDI NPs were measured at each time point. Each bar corresponds with the size results seen in Figure 4.

### Transmission Electron Microscopy

TEM was used to determine the morphology of the NP samples. Figure 6 displays representative images of the empty (A), TMZ-loaded (B), and TMZ+RG7388-loaded (C) double emulsion PS-*b*-PEO+PLGA NPs prepared with functional PLGA in the form of PLGA-NHS and PLGA-NH_2_.

**Figure 6:**
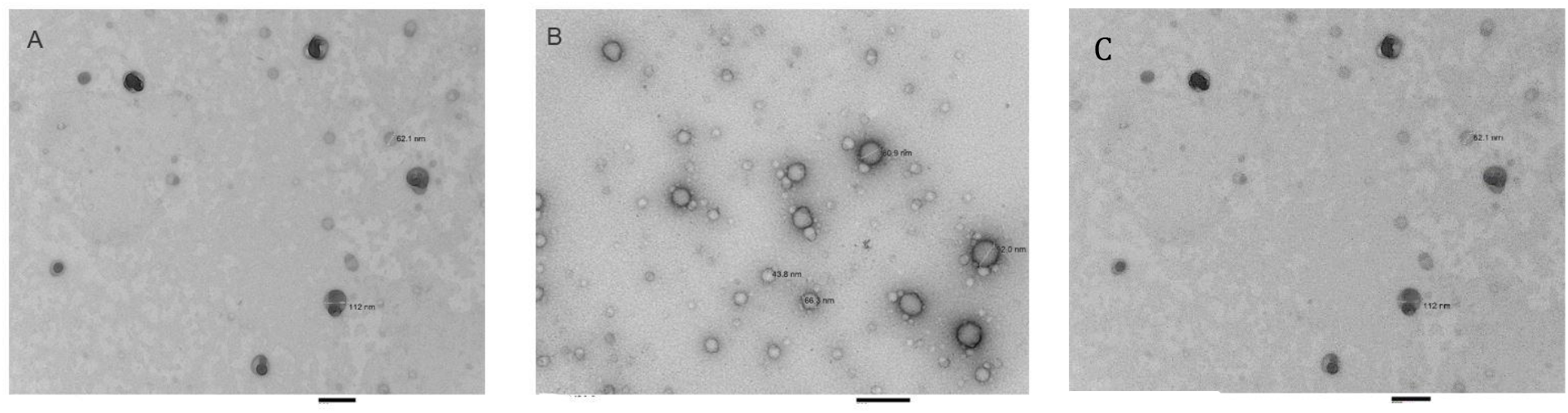
TEM images of control, TMZ-loaded and TMZ+RG7388-loaded NPs. Above are representative TEM images of empty functionalized NPs (A), TMZ-loaded functional NPs, and TMZ+RG7388-loaded NPs (C) fabricated by a double emulsion protocol. Each mag bar correlates to the above image and is 200 nm.

### Drug encapsulation and drug-loading percentage by UV-Vis spectroscopy

Double emulsion NPs containing 50:50 PLGA-NHS/PLGA-NH2 were investigated initially by UV-Vis to see the potential increase in TMZ encapsulation compared to the single emulsion method. The NPs had an EE% of 14.20%±3.23% and a DL% of 1.17%±0.46%.

### Drug encapsulation and drug-loading percentage by HPLC

Encapsulation efficiency and drug-loading percentage were confirmed by HPLC because of the increased sensitivity over UV-Vis spectroscopy. A method to separate TMZ and RG7388 was successfully developed. By HPLC, the encapsulation efficiency and drug-loading percentage of TMZ-loaded NPs was determined to be 4%±6.7% and 0.12%±0.19%, respectively. For TMZ+RG7388-loaded NPs, the encapsulation efficiency was 1.6% for TMZ and 52.8% for RG7388. The drug-loading percentage was 0.07% for TMZ and 0.66% for RG7388.

### Conjugation of CD133 aptamer

The aptamer was bound to the NPs and achieved a conjugation efficiency of 86.3%±7.4%. The EMSA assay was the first method used to determine if the NPs were stably bound to the NPs. According to Figure 7, the free aptamer and aptamer conjugated NPs all shifted approximately the same distance. Once the aptamer was conjugated to TMZ+RG7388-loaded NPs, the size of NPs was 85±3 nm (n=2). In addition, these NPs had a PDI of 0.113±0.011. These final NPs had a zeta-potential of −25.3±0.3 mV. This zeta-potential was significantly more negative than the TMZ+RG7388-loaded NPs without the conjugated aptamer.

**Figure 7.**
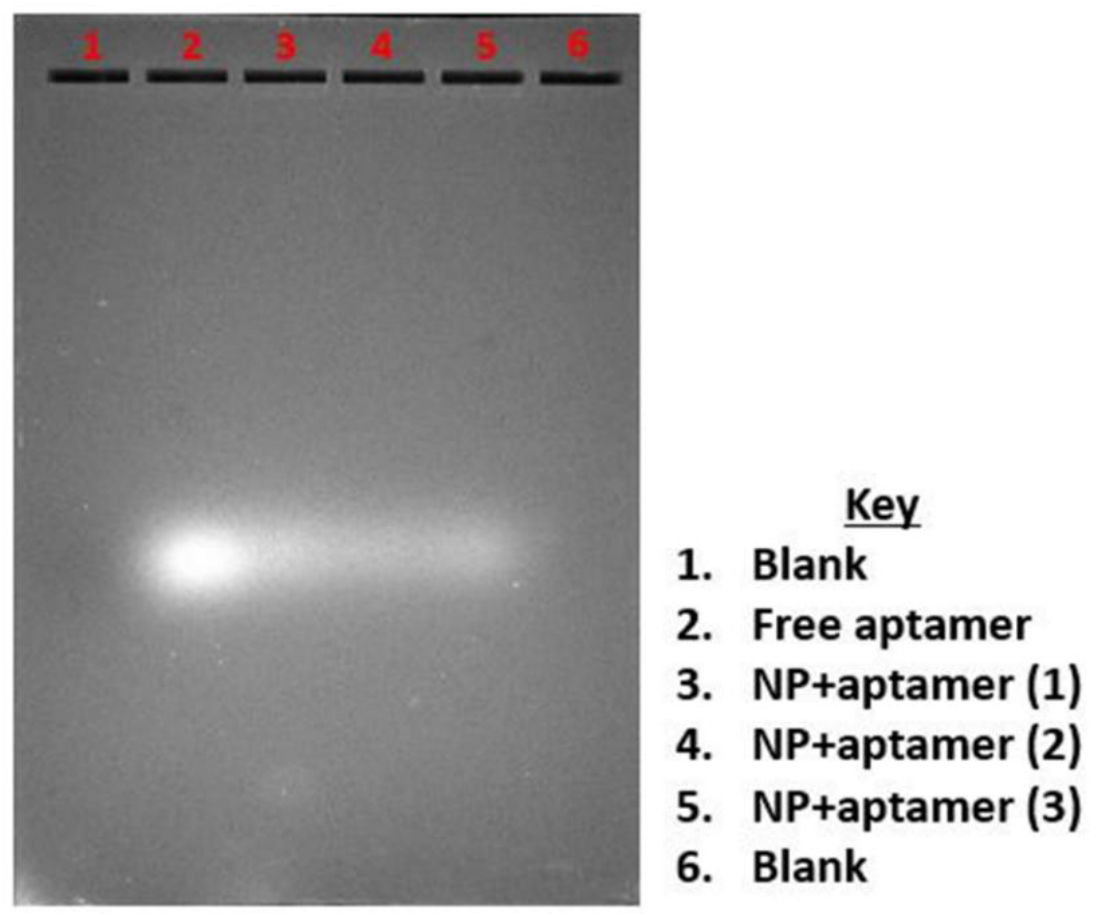
EMSA assay for aptamer bound to NPs. The above figure represents the EMSA assay for NPs bound to aptamers.

The stability of the binding of the aptamer to the NPs was analyzed over a course of 24 hours (Figure 8). The binding efficiency decreased from 94% to 68% within one hour of conjugation. After 3 hours, the binding efficiency was about 50%. From the 3-hour time point to the end of the 24 hours, the binding efficiency increased to 70%.

**Figure 8.**
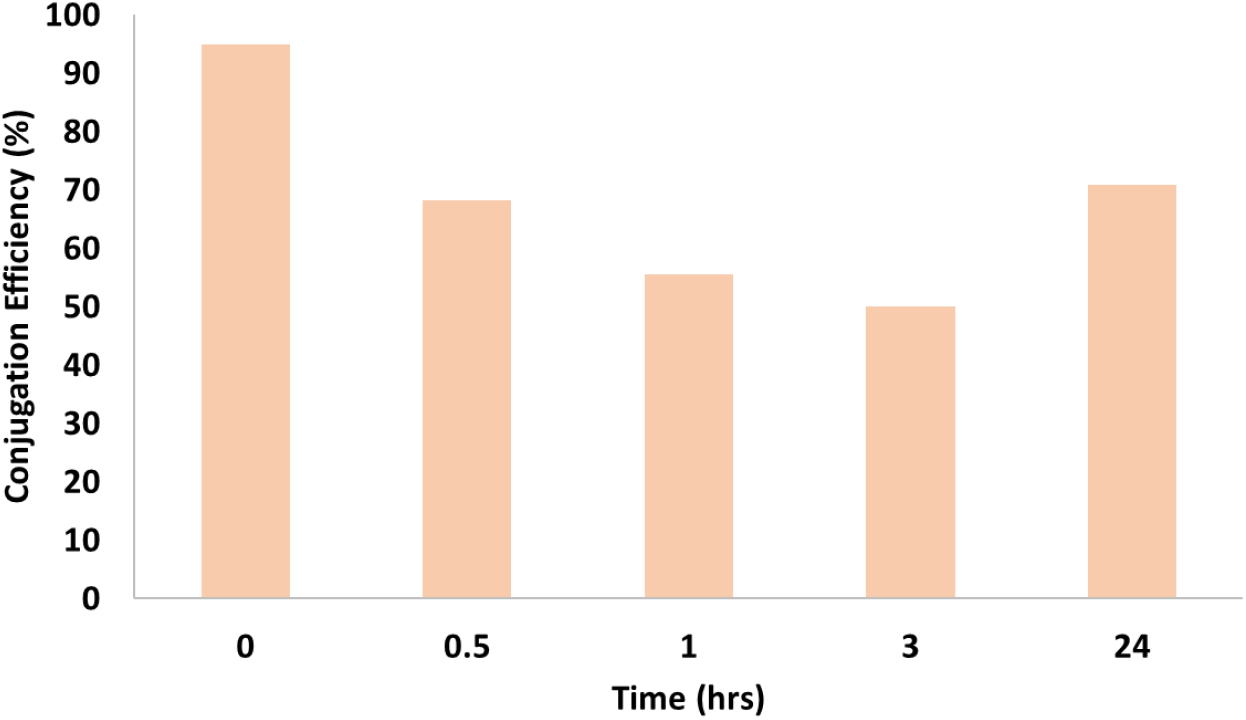
Aptamer stability. The above figure represents the stability of the conjugated aptamer at 37°C over a course of 24 hours.

### Conjugation of ^89^Zr

The first method to conjugated ^89^Zr to the NPs had a labeling efficiency of 29%±2.56% and an average radioactivity of 158.5 μCi. After 24 hours, approximately 46.8%±1.6% of the ^89^Zr released from the NPs. The second method resulted in a significant decrease in binding efficiency to 14%±0.9% with an average radioactivity of 127.7 μCi.

### In vitro analysis

#### Single drug analysis

TMZ and RG7388 single drug analysis was conducted with an n=3 in three different cell populations. Dose response curves can be seen in Figure 9. These Fa values were plotted using a logarithmic analysis to determine the most linear region. These most linear regions were entered into CalcuSyn 2.0 and produced an IC_50_ value for TMZ of 270.02 μM with a lower 95% confidence limit of 214.64 μM and an upper confidence limit of 339.69 μM. The IC_50_ value for RG7388 was determined to be 16.14 μM with a lower 95% confidence limit of 14.58 μM and an upper confidence limit of 17.86 μM.

**Figure 9.**
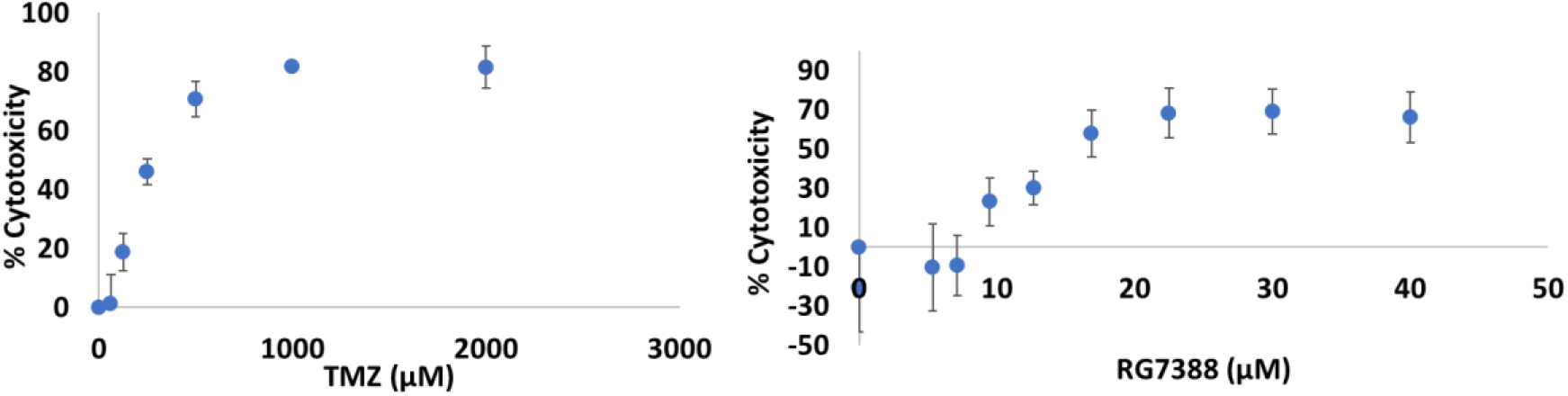
TMZ and RG388 alone in CSCs. The above figure represents the analysis of single drugs in glioma CSCs. The left represents the dose response curve from TMZ and the right represents the dose response curve of RG7388 alone.

A TMZ dose curve was conducted with an n=3 in three cell populations of GBM43 cells. This dose curve can be seen in Figure 10. This was unable to be analyzed via CalcuSyn but produced an approximate IC_50_ value of 338 μM.

**Figure 10.**
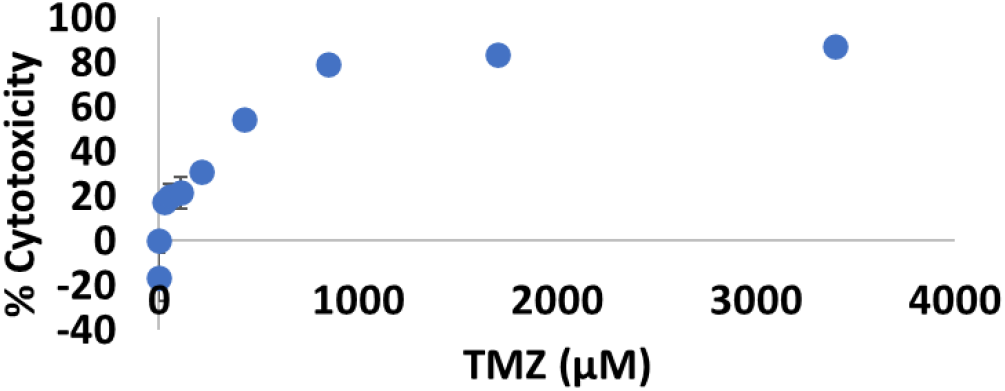
TMZ single drug dose curve in GBM43 cells

#### Combination drug analysis

TMZ and RG7388 were given in combination to CSCs. Ratios of RG7388 to TMZ that were 1:15 and 1:100 were analyzed with an n=3 in three different cell populations. The ratio of 1:50 was only given to n=3 in one population.

These combinations were analyzed using CalcuSyn 2.0 and 50% effective doses (ED50) were determined. These doses were the combination of TMZ and RG7388 that produced 50% killing in CSCs. At a ratio of 15, the ED50 is when TMZ is at a concentration of 122.48 μM. At a ratio of 50, the ED50 is when TMZ is at a concentration of 82.83 μM. Finally, at a ratio of 100, the ED50 is when TMZ is at a concentration of 153.57 μM. The ED50 values were plotted and compared to the effect of the drugs alone. This isobologram in Figure 11 shows an additive effect when RG7388 and TMZ are in a molar ratio of 1:15 and a synergistic effect when the two drugs are in molar ratios of 1:50 and 1:100. This parallels the combination index simulations conducted by CalcuSyn 2.0.

**Figure 11.**
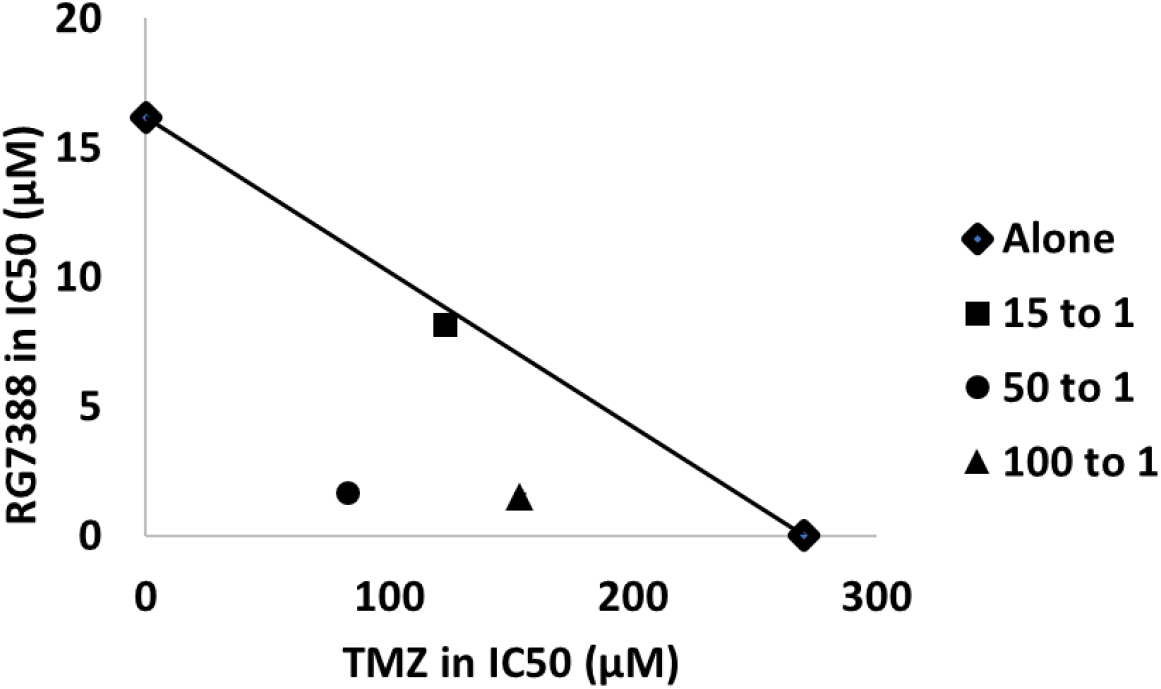
Isobologram of TMZ and RG7388 in CSCs. GBM CSC growth was inhibited when exposed to TMZ in combination with RG7388. RG7388 to TMZ at ratios of 1:50 and 1:100 produced a synergistic effect. Ratio 1:15 produced an additive effect.

### Analysis of treatment with NPs

CSCs received treatments of empty polymer-micellar NPs, TMZ-loaded NPs, TMZ+RG7388 NPs conjugated to the anti-CD133 aptamer. Dose curves for empty NPs and TMZ-loaded NPs are shown in Figure 12. These two sets of NPs had minimal cytotoxic effect on the CSCs. Dose curves for non-targeted (no anti-CD133) dual-drug loaded NPs and targeted (with anti-CD133) dual-drug loaded NPs are shown in Figure 13. These dose curves were indeterminate using CalcuSyn 2.0. However, a similar analysis was conducted to determine an approximate IC50 value. Dual drug-loaded NPs had an IC50 value of the TMZ concentration of 14.26 μM. The addition of the targeting agent somewhat lowered the IC50 value to a TMZ concentration of 11.86 μM.

**Figure 12.**
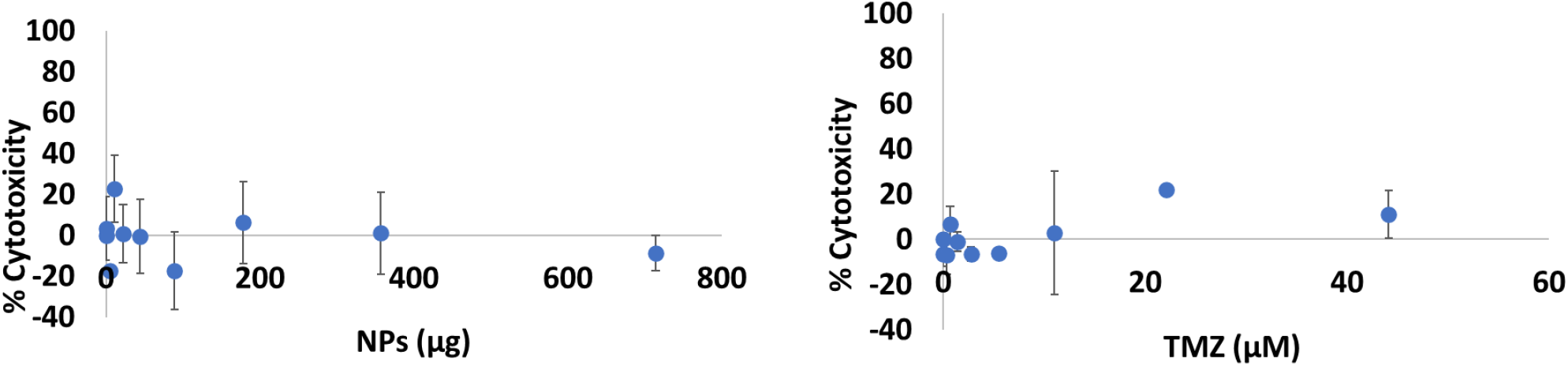
Empty and TMZ-loaded NPs in CSCs. Above represents the dose curves generated from empty polymer-micellar NPs (left) and TMZ-loaded NPs (right). The mass of NPs is the same in each dose curve.

**Figure 13.**
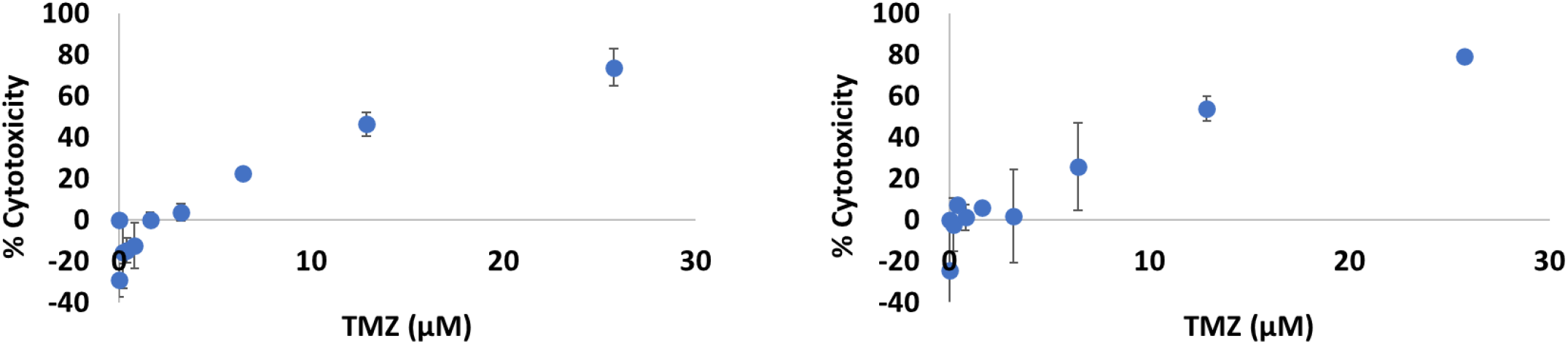
Non-targeted TMZ+RG7388-loaded NPs and targeted TMZ+RG7388-loaded NPs in CSCs. Above represents the dose curves generated from dual-drug NPs (left) and dual-drug NPs conjugated to the anti-CD133 aptamer (right).

## Discussion

The quest for a cure for GBM has been disappointing since no new drug has significantly impacted patient survival in more than fifteen years. A single drug approach to disease treatment is unlikely to be a viable option given the high propensity of GBM for recurrence, which is at least partially driven by the highly resistant subpopulation of self-regenerating GBM CSCs (7, 8). NPs hold great promise for overcoming the limitations of a single drug approach by permitting multi-drug combinations that treat GBM synergistically. In addition, NPs open the door to utilizing the unique advantages of theranostics since the therapeutic agent can be assessed using an imaging label. The goal of these experiments was to establish the building blocks of a multi-drug therapeutic approach that could systematically dismantle GBM by preferentially targeting and overriding the intrinsic TMZ resistance of GBM CSCs. In addition, an integrated imaging label was sought that would permit a fully theranostic approach to GBM. For this work, four hypotheses were generated: 1.) Hybrid functionalized PS-*b*-PEO and PLGA NPs can be harnessed to encapsulate two different chemotherapies, TMZ and RG7388, to kill human-derived GBM CSCs in a dose dependent manner in vitro. 2.) A 15 nucleoside CD133 aptamer can be covalently bound to dual-drug loaded PS-*b*-PEO-PLGA NPs to selectively target the CD133 antigen on the CSC surface. 3.) Chelative binding of the ^89^Zr radiotracer to the surface of the NPs would establish a theranostics application for in vivo PET biodistribution studies. 4.) An NP size of less than 100 nm and acceptable polydispersity could be achieved for the final dual-drug loaded, functionalized NPs.

Steady progress was made towards the first hypothesis by constructing PS-*b*-PEO and PLGA NPs that encapsulated both TMZ and RG7388. The original goal had been to maintain a size of 25-50 nm consistently published by collaborators from Ohio State (Jessica Winters, Ph.D.) (28).

A size near 50 nm was achieved, but only during the production of NPs without drugs and without functional groups. As more components were added to the NPs in a stepwise fashion, the NPs increased in size above 50 nm, but the final NPs with two drugs and two functional groups stayed below 100 nm. In contrast, dual drug-loaded mPEG-PLA NPs also encapsulating two drugs (TMZ and PTX) were of a 200 nm size (29). It is possible that a hybrid approach using both micelles and polymer improved the size of the NPs compared with traditional polymer PLGA particles (28, 30). In the process of optimizing drug encapsulation, a double emulsion technique modified from Xu et al., permitted dual-encapsulation with both TMZ and RG7388 more efficiently than a single emulsification as reported in the literature (26, 28). As shown in the results, size of dual drug-loaded NPs was reduced from 260 nm down to 85 nm using this technique. Other published studies utilized sizes of NPs of 275 nm, 300-500 nm, and 206 nm (26, 29, 31, 32). These results may have been influenced by use of polymer components consisting of PLGA-PEG-PLGA, PCL, and PLGA-mPEG (29, 31, 32). While NPs fabricated by a double emulsification technique typically result in larger sizes, others have achieved a size as small as 40 nm when co-loading 5-fluorouracil and Chrysin into PLGA-PEG-PLGA NPs (33). During the production of NPs, a PDI of 20% or less was maintained which is desirable since lower polydispersity indicates a more uniform size distribution (34). As for charge, a slightly negative charge was achieved for dual drug-loaded particles, except during addition of the CD133 aptamer, which decreased the zeta potential from −8 mV to −25 mV. While the ideal charge is not known, there is a general consensus that a neutral or slightly negative charge neither repulses the cell membrane nor binds it too tightly (35). Finally, dual-drug PS-*b*-PEO+PLGA NPs exhibited a consistent spherical morphology when analyzed by TEM and revealed a low level of clumping even with longer duration in solution. Particles with a more spherical shape can potentially result in better cell uptake compared to cylindrical particles as well as increased circulation time (34).

With successful construction of TMZ+RG7388 NPs, NPs were compared with blank control NPs and NPs containing only TMZ. synergistic killing of CSCs was first demonstrated using combination therapy of TMZ and RG7388, through an analysis of the CI outlined by the Chou-Talalay method, without using NPs first. Through these results, a reduction in the IC_50_ value of TMZ from 270 μM to 83 μM was achieved in combination with RG7388 at a molar ratio of 50:1. A prior published study from a synergistic effect between TMZ and RG7388 in the human derived GBM10 cell line, but to our knowledge, the GBM CSC population has not been treated with this drug combination (17). Dual drug-loaded NPs produced a higher killing effect than empty NPs and TMZ-loaded NPs to achieve the fifth objective.

Following initial cell killing studies with non-targeted NPs, the second hypothesis was addressed to develop the CD133 aptamer binding to the TMZ+RG7388 NPs. A high conjugation efficiency of 86% was achieved with the fluorescently labelled CD133. This accomplished the third objective. An attempt was made to validate the stability results using an EMSA. Although the NPs are much larger in size compared to the aptamer, charge also influences the degree to which the compounds will migrate. Both free aptamer and aptamer bound to the NPs travelled the same distance approximately 60% from the starting point creating a single well-defined band without a gradient in the EMSA study.

Dissociation of CD133 aptamer from the NPs would produce a non-uniform gradient. A single preliminary study of the stability of CD133 aptamer binding to the NPs revealed 50% aptamer remained bound to the NPs at 3 hours and 60% still bound at 24 hours. These results are difficult to interpret since the experiment was performed only once and needs to be replicated in future trials.

To address the second hypothesis, superior CSC killing was attempted with TMZ+RG7388+CD133 NPs compared with TMZ+RG7388 NPs without CD133. There was a trend towards a lower IC_50_ value in CD133 aptamer-bound NPs compared to those without the CD133 aptamer. Future in vitro and in vivo studies need to be conducted to assess the potential advantages of the CD133-labelled NPs.

To address the third hypothesis, NPs were labeled with ^89^Zr using two different methods over three trials. In the first method NPs were incubated in a DFOM solution for 40 minutes and labelled with ^89^Zr for one hour to achieve 30% labelling. To try to improve labeling, the NPs were incubated in the DFOM solution overnight and labelled with the ^89^Zr for two hours. The first method achieved a higher amount of ^89^Zr labelled to the NPs and had a final radioactivity of 158 μCi on 1.5 mg of NPs. For future in vivo studies, this technique will need to be optimized to meet the 250 μCi minimum in each dose of NPs for our current Indiana University IndyPET preclinical system. These initial steps will help us further achieve the fourth objective.

Overall encapsulation efficiency of TMZ in the NPs was low but was higher than reported in the literature. For instance, one study extensively analyzed a variety of ways to improve the encapsulation of TMZ and achieved a maximum of 2% encapsulation (26). To overcome this challenge, a series of studies were conducted to understand the stability and optimal environments for using TMZ in the development of NPs. After several different techniques were applied, TMZ encapsulation remained low with minimal killing of GBM CSCs following treatment with TMZ-loaded NPs. However, encapsulation of TMZ in combination with RG7388 likely still contributed to combination therapy based on the in vitro results. Future studies could test delivery of TMZ orally as part of standard therapy, and then deliver two different drugs more feasibly encapsulated into the NPs. Such a system would allow a three-drug approach to supplement oral TMZ therapy.

Another barrier to successful TMZ encapsulation was lack of uniformity across the current published literature with regard to improved formulations of NPs or measurement techniques. While some publications report encapsulation efficiencies using a direct measurement of drug content in a pellet of NPs, others use indirect measurements that may result in an apparently higher, although less accurate encapsulation efficiency (29). To improve data accuracy and minimize unnecessary and repetitive formulation changes, the nanotechnology community has recently proposed more standardized protocols for data reporting (36).

### Limitations

There were limitations encountered which would ideally be addressed in future work. The most important and time-consuming limitation was encapsulation of TMZ. Options would be to either replace TMZ with a suitable anticancer drug or try a different encapsulation approach. For instance, a w/o/o phase separation in a non-aqueous medium could be studied. In this method, both hydrophilic and lipophilic drugs can be encapsulated while avoiding the possibility that aqueous solutions wash away hydrophilic drugs (37). Instead of emulsifying the components in a larger volume of aqueous solution, the hydrophilic drug is mixed into an organic solvent in the presence of an organic nonsolvent, such as silicone oil, that will not dissolve either drug or the polymers (37). As the organic solvent evaporates, this forces the polymers to undergo a phase separation and the polymer will adsorb onto the drug (37). This method may increase the encapsulation of TMZ but result in a size increase of the NPs.

If TMZ was eliminated from the particle system and used as an oral agent, new dual-drug NPs could be developed that do not cross the BBB as well as orally delivered TMZ. Given the much higher potency of RG7388 encountered in treating CSCs, TMZ encapsulation may need to increase significantly for a combination effect to be recordable. While studying drug effects on CSCs in the absence of NPs, much higher concentrations of TMZ were utilized for comparison with RG7388. Higher amounts of RG7388 and low TMZ may result in predominately RG7388-induced toxicity RG7388 could be encapsulated with another potent hydrophobic drug such as PTX that offers better encapsulation efficiency and greater cell killing.

Additional studies to show the potential benefit of CD133-targeted NPs need to be conducted. The current results show a slight increase in cytotoxicity with the addition of the CD133 aptamer to the NPs. However, because this increase had not been yet shown to be a significant change, the drugs may be releasing into the wells regardless of whether the NPs were bound to the CSCs. First, additional cytotoxicity trials are needed. Next, to better assess the targeting ability of CD133-labelled NPs two different fluorescent techniques could be attempted. The first option would be to assess uptake of CD133 targeted and non-CD133 targeted NPs into cells using fluorescence correlation spectroscopy (FCS) which assesses whether particles are taken into cells in vitro by the presence of a diffusion coefficient measured by the fluorescent marker. Initial studies were conducted to determine the diffusion coefficient of NPs bound to the fluorescent aptamer by suspending them in water. However, during these studies, the NPs produced a flickering effect, which introduced inconsistencies in the correlation analysis. Once the diffusion coefficient is determined, the aptamer-conjugated NPs can be administered to populations of CSCs. 24-well or 6-well plates would increase the number of CSCs for assessment compared with the 96 well plates used in this study. CSCs could be treated for up 24 hours with the aptamer-conjugated NPs, fixed with paraformaldehyde and then analyzed using FCS. If the CSCs take up the NPs bound to the aptamer, an increase in the diffusion coefficient would be visible. To compare targeted verses non-targeted NPs, either the PLGA or the PS*b*-PEO could be labeled with a fluorescent dye such as Cy5 dye for assessing whether more CSCs take up the targeted or non-targeted CSCs. Second, fluorescent imaging microscopy could be performed permitting image acquisition of the Cy5 labelled NPs to assess cell uptake of targeted verses non-targeted NPs. Images could also be taken analyzing the uptake ability of CD133+ CSCs and the CD133-CSCs. To do this, the CSC population would first need to be sorted using fluorescence-activated cell sorting (FACS) by immunostaining the CD133+ population prior to sorting. This study could also be conducted in co-culture with CD133+ CSCs and neuronal stem cells that do not express CD133. This would allow assessment of non-specific targeting. Finally, differential killing of CD133 and non-CD133 labeled NPs may ultimately be further assessed in vivo.

Future improvements of the labelling of ^89^Zr also need to be made. In vivo studies cannot be performed until more ^89^Zr is tagged to the NP surface. One way to improve the labelling efficiency could be using an alternate chelating agent. Previous work by Veronesi et al. used a derivative of DFO that was attached to a benzyl isothiocyanate (DFO-Bz-NCS) (38). Furthermore, another study showed an increased in binding and stability using DFO* as part of the above derivative which adds an additional hydroxamic acid function to the chelate chain compared to DFO alone (39).

In summary, important progress was made in vitro towards the formation and initial testing of a multifunctional theranostics nanoparticle that can be used to target and treat the CSC specific cell population with GBM. This work comprised a significant portion of the in vitro arm of the project. The second arm of the project will be to test the NP compound in vivo in a preclinical model of GBM for both therapy efficacy and fate mapping.

## CONCLUSION

GBM is a devastating disease with a dismal prognosis and a paucity of effective chemotherapeutic agents. The increased mutation rate, infiltrative nature and high propensity for self-renewal of the CSC population are important reasons for failure of single drug regimens against GBM. This work outlines a strategy for using multi-functional polymer-micellar NPs to target the CSC population and deliver a therapy that overcomes GBM resistance to TMZ. PLGA-NHS and PLGA-NH2 functional groups were utilized as co-linkers to attach a CD133 aptamer and radiotracer to the NPs while also maintaining a relatively uniform size of less than 100 nm with a slightly negative charge. Preliminary CD133 labeling of NPs demonstrated a trend in higher killing of CSCs in vitro compared with CD133 deficient NPs. Addition of PET radiotracer, ^89^Zr, to the multi-functional CD133 targetable NPs introduced a potential theranostics application to allow real time in vivo fate mapping.

## Acknowledgements

Dr. Chien-Chi Lin and Dr. Mangilal Agarwal

Integrated Nanosystems Development Institute (INDI)

Barbara Bailey and the Indiana University Simon Cancer Center In Vivo Therapeutics Core Dr. Mosa Alhamami

Julia Payton

Sherry Clemens

## APPENDIX

## Materials

### Nanoparticle fabrication

Carboxyl-terminated PS-*b*-PEO (molecular weight 5,000-*b*-2,200, Cat No P4090SEOCOOH) and carboxyl-terminated PS-*b*-PEO (molecular weight 9,500-*b*-18000 Da, Cat No P18154-SEOCOOH, Polymer Source Inc., Montreal, QC, Canada). PLGA (molecular wieght 73,000 Da, Cat No APO60), PLGANH2 (diamine) (PLGA-NH2) (molecular weight 30,000-40,000 Da, Cat No AI062) and PLGA-N-hydroxysuccinimide (PLGA-NHS) (molecular weight 50,000-80,000 Da, Cat No AI116 PolySciTech, West Lafayette, IN). Poly(vinyl alcohol) (PVA) (molecular weight 13,000-23,000 Da, Cat No 348406) and PVA (molecular weight 31000-50000, Cat No 363138, Sigma Aldrich, St. Louis, MO). Pluoronic F-68 Prill 188 (Material 30085243, BASF, Mount Olive, NJ). Temozolomide (TMZ) (Cat No T2577, Sigma Aldrich, St. Louis, MO). RG7388 (Cat NO HY-15676/Cs-1473, MedChem Express, Monmouth Junction, NJ). Hydrochloric acid (HCl) (0.1N, Cat No S25354), Dichloromethane (DCM) (Cat No AC406920010, Fisher Scientific, Waltham, MA). Formic acid (≤95%, Cat No F0507-500mL, Sigma Aldrich, St. Louis, MO). Acetonitrile (ACN) (Cat No A996-4, Fisher Scientific, Waltham, MA). Ethyl acetate (Cat No 035909, Oakwood Chemical, Estill, SC).

### Nanoparticle conjugation

Non-fluorescent CD133 aptamer (5’(C6-NH2) CCC UCC UAC AUA GGG 3’ PO RNA) (Cat No O-5100) was purchased from TriLink Biotechnologies (San Diego, CA).

Fluorescein amidite (FAM)-azide labelled CD133 aptamer (5’ C6-NH2) CCC UCC UAC AUA GGG (FAM-Azide) 3’) was purchased from Integrated DNA Technologies, INC. Water (for RNA work) (Cat No BP561-1) was purchased from Fisher Scientific (Waltham, MA). Tris-Acetate-EDTA (TAE) (10X solution, Cat No BP1335-1) was purchased from Fisher Scientific (Waltham, MA). ^89^Zr(HPO_4_)_2_ solution was from Washington University (St. Louis, MO). Deferoxamine mesylate salt, the mesylate salt of DFO, (DFOM) (≤92.5% (TLC) Cat No D9533) was purchased from Sigma Aldrich (St. Louis, MO).

### Polymers/Surfactant

Carboxylic acid terminated poly(styrene-*b-*ethylene oxide) (PS-*b*-PEO) (molecular weight 9500-b-18000 Da, Cat No P18154-SEOCOOH) was purchased from Polymer Source Inc. (Montreal, QC, Canada). Poly(lactide-*co*-glycolide) (PLGA) (molecular weight 73,000 Da, Cat No AP060), poly(lactide-*co*-glycolide)-NH_2_ (diamine) (PLGA-NH_2_) (molecular weight 30,000-40,000 Da, Cat No AI062), and poly(lactide-*co*-glycolide-N-hydroxysuccinimide (PLGA-NHS) (molecular weight 50,000-80,000 Da, Cat No AI116) were purchased from PolySciTech (West Lafayette, IN). Poly(vinyl alcohol) (PVA) (molecular weight 13,000-23000 Da(Cat No 348406, Sigma Aldrich, St. Louis, MO).

### Drugs/Aptamers

Temozolomide (TMZ) (Cat No T2577) was purchased from Sigma Aldrich (St. Louis, MO). RG7388 (Cat No HY-15676/Cs-1473) was purchased from MedChem Express (Monmouth Junction, NJ). CD133 (5’ (C6-NH_2_) CCC UCC UAC AUA GGG 3’ PO RNA) (Cat No O-5100) was purchased from TriLink Biotechnologies (San Diego, CA). PE labelled anti-human CD133 antibody, clone 7 (Cat No 372803) supplied by BioLegend (San Diego, CA).

### Cell studies

Human GBM43 cells were kindly donated from the Simon Cancer Center at Indiana University. Culture media was Gibco Dulbecco’s Modified Eagle Medium (DMEM) with 4.5 g/L D-Glucose, L-Glutamine, Life-Technologies, Grand Island, NY) with 10% Fetal bovine serum (FBS) (Cat No 35-016-CV, Corning Inc., Corning, NY) and 1% HEPES buffer (1M pH 7.3, Cat No 118-089-721, Quality Biological, Gaithersburg, MD). Methylene blue (1% in ethanol, Cat No LC169201, LabChem (Zelienople, PA). Human GBM cancer stem cells (CSCs) (Cat No 36104-41, Celprogen, Torrance, CA). Culture media was Human Glioma Cancer Stem Cells Media with Serum (Cat No M36104-40S without antibiotics, Celprogen, Torrance, CA). Methylene blue (1% in ethanol, Cat No LC169201, LabChem, Zelienople, PA)

